# Type I IFN signaling in the absence of IRGM1 promotes *M. tuberculosis* replication in immune cells by suppressing T cell responses

**DOI:** 10.1101/2023.10.03.560720

**Authors:** Sumanta K. Naik, Michael E. McNehlan, Yassin Mreyoud, Rachel L. Kinsella, Asya Smirnov, Chanchal Sur Chowdhury, Samuel R. McKee, Neha Dubey, Reilly Woodson, Darren Kreamalmeyer, Christina L. Stallings

**Affiliations:** Department of Molecular Microbiology, Center for Women’s Infectious Disease Research, Washington University School of Medicine, St. Louis, MO 63110, USA

**Keywords:** *Mycobacterium tuberculosis*, macrophage, neutrophil, IRGM1, IRGM3, inflammation, T cell, type I interferon, IFN-γ, lung, lymph node

## Abstract

Polymorphisms in the *IRGM* gene are associated with susceptibility to tuberculosis in humans. A murine ortholog of *Irgm*, *Irgm1*, is also essential for controlling *Mycobacterium tuberculosis* (Mtb) infection in mice. Multiple processes have been associated with IRGM1 activity that could impact the host response to Mtb infection, including roles in autophagy-mediated pathogen clearance and expansion of activated T cells. However, what IRGM1-mediated pathway is necessary to control Mtb infection *in vivo* and the mechanistic basis for this control remains unknown. We dissected the contribution of IRGM1 to immune control of Mtb pathogenesis *in vivo* and found that *Irgm1* deletion leads to higher levels of IRGM3-dependent type I interferon signaling. The increased type I interferon signaling precludes T cell expansion during Mtb infection. The absence of Mtb-specific T cell expansion in *Irgm1*^-/-^ mice results in uncontrolled Mtb infection in neutrophils and alveolar macrophages, which directly contributes to susceptibility to infection. Together, our studies reveal that IRGM1 is required to promote T cell-mediated control of Mtb infection in neutrophils, which is essential for the survival of Mtb-infected mice. These studies also uncover new ways type I interferon signaling can impact T_H_1 immune responses.

## INTRODUCTION

*Mycobacterium tuberculosis* (Mtb) infection is a leading cause of death world-wide. The COVID-19 pandemic has reversed years of global efforts to tackle the tuberculosis (TB) pandemic, where for the first time in over a decade, TB deaths have increased over the last year^1^. The clinical outcomes of TB are driven by the immune response mounted in the host. Because of the pivotal role of the immune response in TB, there is a growing interest in developing immunotherapies that harness the immune response to control the infection, however, this will require a better understanding of the immune responses required to control Mtb infection. Interferon (IFN) inducible immunity-related GTPases (IRGs) play important roles in the host immune response to intracellular pathogens^2^, where a polymorphism in the *IRGM* gene that decreases its expression has been associated with Crohn’s disease and active TB disease in humans^3,4^. Mice encode three IRGM orthologs, IRGM1, IRGM2, IRGM3, that have been shown to have non-redundant functions. In particular, *Irgm1*^-/-^ mice infected with Mtb succumb early with higher bacterial burdens and larger lesions of inflammation in their lungs^5,6^, supporting a requirement for IRGM1 expression for a protective immune response to Mtb. Deletion of *Irgm3* rescues the higher bacterial burdens and early susceptibility in *Irgm1*^-/-^ mice during Mtb infection^5^, indicating that a balance of IRGM1 and IRGM3 is required to control Mtb pathogenesis. However, the underlying mechanism for why IRGM1 is required in the presence of IRGM3 expression to control Mtb infection remains unknown.

Inter-regulatory roles for IRGM homologs have been implicated in multiple processes, where the most extensively studied has been the role for IRGM orthologs in autophagy. In particular, decreased expression of IRGM1 results in defects in autophagy in epithelial cells and phagocytes *in vitro*^3,7–9^, which can be reversed by deletion of *Irgm3*^10,11^. In initial studies, IRGM1 was shown to co-localize with Mtb in autophagosomes within infected macrophages and facilitate the trafficking of Mtb to lysosomes *in vitro*^6,12–14^, which supported an original model that IRGM1 controls Mtb replication by promoting autophagy-mediated clearance of the pathogen. However, more recently this was drawn into question by a report that IRGM1 does not colocalize with phagosomes containing *Mycobacterium bovis* BCG in IFN-γ-treated cells^15^. In addition, we have previously shown that autophagy is not required within infected myeloid cells to control Mtb replication *in vivo* in the mouse models used to study *Irgm1*^16,17^. Therefore, defects in autophagy-mediated targeting of Mtb to lysosomes within infected cells is unlikely to explain the susceptibility of *Irgm1*^-/-^ mice to Mtb.

More recently, studies investigating the role for *Irgm1* during mycobacterial infection showed that *Irgm1*^-/-^ mice are unable to control *Mycobacterium avium* replication and succumb to the infection earlier than wild-type (WT) controls due to IFN-γ-induced autophagic cell death of T cells following priming and activation^18^. *Irgm1*-deficient T cells were able to rescue the susceptibility of *RAG2*^-/-^ mice to *M. avium* infection, suggesting that the defect in survival following activation was not T cell intrinsic^18^. However, *in vitro* studies revealed that IRGM1-deficient T cells exhibit decreased survival and defective expansion following activation in the presence of IFN-γ, suggesting that there are T cell intrinsic roles for IRGM1 as well as roles for IRGM1 in other cells that contribute to survival of T cells following activation in the presence of IFN-γ^19^. In addition to the effects on T cells following activation, *Irgm1*^-/-^ mice also exhibit IFN-γ-dependent hematopoietic stem cell (HSC) hyperproliferation and defects in HSC differentiation during *M. avium* infection, which may also contribute to the lymphopenia^20,21^. Pulmonary Mtb infection of *Irgm1*^-/-^ mice was also accompanied by decreased cellularity and lower T cell numbers in the draining lymph node (dLN) compared to WT controls^18^. Similar observations have been reported during infection with the intestinal pathogen *Citrobacter rodentium*, where *Irgm1*^-/-^ mice exhibit a defect in T_H_1 cell expansion and myeloid cell survival following infection^22^. The decreased T_H_1 cell expansion and myeloid cell survival following *C. rodentium* infection was attributed to increased apoptosis in the absence of IRGM1 expression. Macrophages from *Irgm1*^-/-^ mice infected with *Salmonella typhimurium* also exhibit impaired IFN-γ responses, including defects in adhesion, motility, and killing of the bacteria^23^. *Irgm3* deletion reverses the effects of *Irgm1* deletion on IFN-γ responses in HSCs^20,23^, but whether *Irgm3* deletion would reverse hypersensitivity to IFN-γ in other cell types has yet to be determined.

These prior studies highlight multiple potential roles for IRGM1 that could affect control of Mtb infection, however, no specific role has been directly shown to contribute to the susceptibility. In this manuscript, we dissect the mechanistic basis for IRGM1-mediated protection against Mtb infection *in vivo*. We found that *Irgm1* deletion leads to higher levels of IRGM3-dependent type I IFN signaling in naïve mice. The increased type I IFN conditions an environment where following Mtb infection, dendritic cells (DCs) are less abundant in the lungs and dLN, and the T cell population is unable to expand following activation in the dLN. The absence of Mtb-specific T cell responses results in uncontrolled Mtb infection in neutrophils and alveolar macrophages (AMs) within the lung, which directly contributes to susceptibility to Mtb infection. Therefore, our data not only reveal the basis for IRGM1-mediated protection during Mtb infection *in vivo*, they also reveal a previously unknown consequence of increased type I IFN signaling on T_H_1 responses, which could in part explain the association of type I IFN with poor TB disease outcomes in humans and mice and could also apply to other infections requiring T_H_1 immunity for control.

## RESULTS

### Susceptibility of *Irgm1*^-/-^ mice to Mtb infection is associated with impaired T cell expansion in the dLN and defects in innate immune cells in the lung

To investigate the mechanistic basis for the requirement of IRGM1 in protection against Mtb infection, we used a mouse model of pulmonary Mtb infection. Following aerosol infection of a WT mouse, alveolar macrophages (AMs) phagocytose Mtb and traffic to the parenchyma, spreading the bacteria to other immune cells^24–26^. Monocyte-derived DCs transport Mtb antigen to the dLN at 8-9 days post infection (dpi) and facilitate priming, activation, and proliferation of Mtb-specific T cells^27–29^. Effector T cells then traffic to the lungs beginning at 18 dpi and activate the infected innate immune cells to suppress Mtb replication^30^, allowing for survival of a WT mouse for over 150 days. Infection of *Irgm1^-/-^*mice with an Mtb strain that constitutively expresses GFP^31,32^ (Mtb-GFP) results in early susceptibility (median survival of 41 days) **(Fig. 1A)** and increased lung burden at 21 dpi **(Fig. 1B)**, similar to as previously reported^5,6^. Consistent with prior studies^19^, we also observed a defect in T cell expansion in the dLN of Mtb-infected *Irgm1*^-/-^ mice, as evidenced by smaller lymph nodes size **(Fig. 1C)** and decreased T cell numbers in the dLN at 21 dpi **(Fig. 1D)**. The lower numbers of T cells were associated with a lower number of activated CD4^+^ T cells (CD44^+^CD62L^-^) in the dLN **(Fig. 1E)**. We also observed lower numbers of DCs in the dLN of *Irgm1*^-/-^ mice compared to controls at 21 dpi **(Fig. 1F)**, which could contribute to the absence of T cell expansion following infection. In addition, we monitored T cell viability using the Zombie Viability Dye (BioLegend) and observed decreased viability in activated CD4^+^ T cells in the dLN of *Irgm1*^-/-^ mice compared to controls at 21 dpi **(Fig. 1G)**, which supports the prior model that IRGM1 is required for T cell survival following activation.

**Fig. 1:**
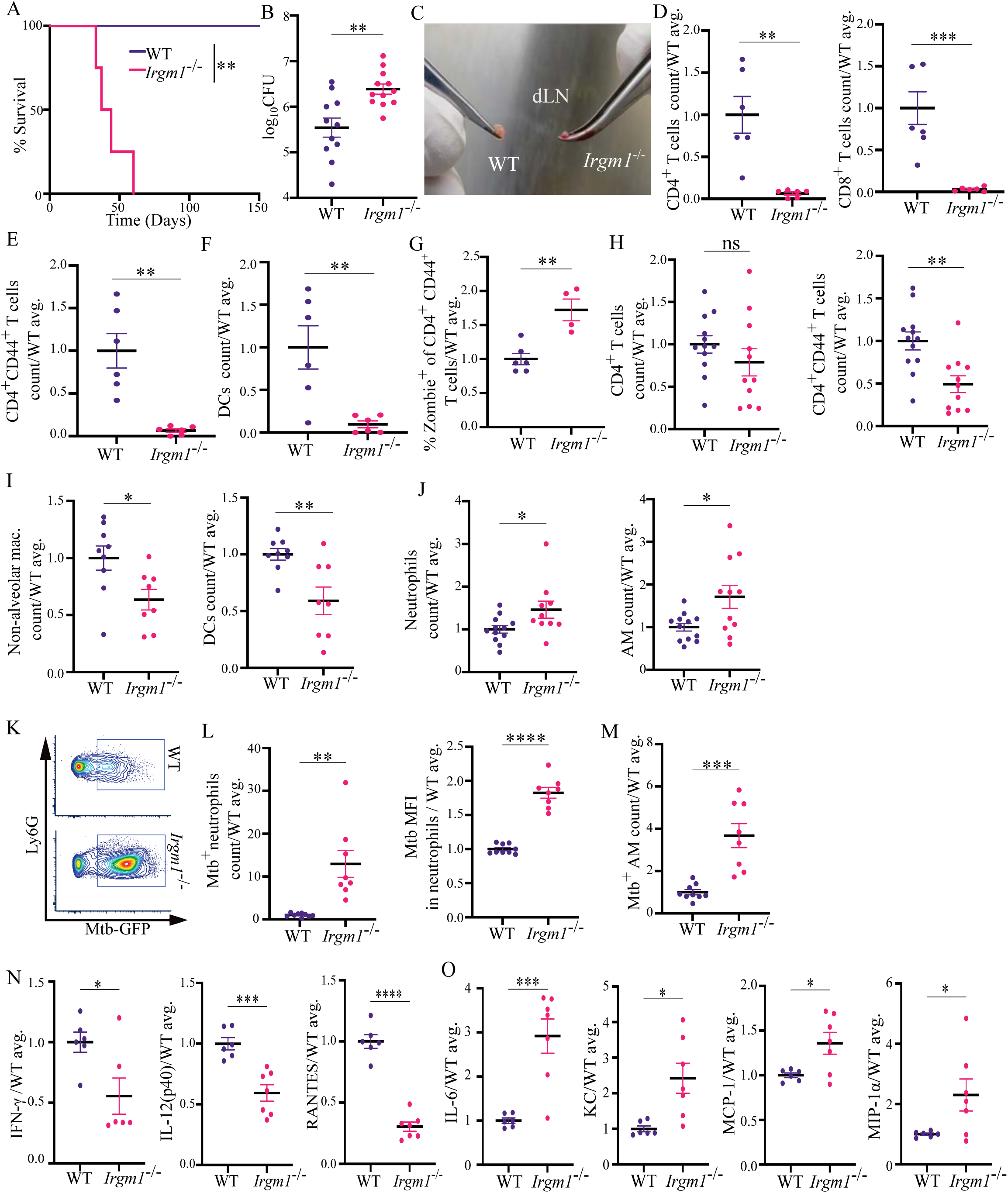
Susceptibility of *Irgm1*^-/-^ mice to Mtb infection is associated with impaired T cell expansion in the dLN and defects in innate immune cells in the lung (A) Survival of WT and *Irgm1^-/-^* mice following Mtb aerosol infection (WT n=5, *Irgm1*^-/-^ n=4). Statistical significance of differences was determined by the Mantel-Cox test (**P < 0.01). **(B)** Mtb CFU count in lungs at 21 dpi. **(C)** Representative mediastinal dLN from WT and *Irgm1^-/-^* mice at 21 dpi. **(D-F)** Number of **(D)** CD4^+^ and CD8^+^ T cells, **(E)** activated CD4^+^ T cells (CD44^+^CD62L^-^), and **(F)** DCs in the dLN at 21 dpi. **(G)** % of activated CD4^+^ T cells that were Zombie^+^ (nonviable) in the dLN at 21 dpi. **(H-J)** Number of **(H)** CD4^+^ T cells and activated CD4^+^ T cells, **(I)** non-alveolar macrophages and DCs, and **(J)** neutrophils and alveolar macrophages (AM) in the lungs at 21 dpi. **(K)** Representative flow plot showing Mtb-GFP positivity in Ly6G^+^ neutrophils in the lung at 21 dpi. **(L)** Number of Mtb^+^ (based on GFP from Mtb-GFP) neutrophils and Mtb burden (based on mean fluorescence intensity (MFI) of GFP) per infected neutrophil in the lung at 21 dpi. **(M)** Number of Mtb^+^ AMs in the lung at 21 dpi. **(N-O)** Cytokine and chemokine levels in bulk lung homogenate at 21 dpi. Each data point corresponds to one mouse and graphs show the mean ± SEM, where in **(D-J, L-O)** the data are expressed as the ratio to the average for WT in a single infection experiment to mitigate inter-experiment variation in total cell counts. **(D-J)** Data is combined from experiments using both WT Mtb Erdman strain and GFP expressing Mtb (GFP-Mtb). **(B, D-J, L-O)** Statistical significance of differences were determined by two-tailed unpaired t-tests. **** P < 0.0001; *** P < 0.001; ** P < 0.01; * P < 0.05; ns=not significant.

To determine how immune responses in the lung were affected by the defective dLN responses in *Irgm1*^-/-^ mice during Mtb infection, we performed flow cytometry analysis of immune cell populations in the lungs at 21 dpi. We observed decreased numbers of activated CD4^+^ T cells in the lungs of *Irgm1^-/-^* mice compared to WT mice at 21 dpi **(Fig. 1H)**, which is likely a result of the lack of T cell expansion in the dLN of Mtb-infected *Irgm1*^-/-^ mice **(Fig. 1C-E)**. In addition, the numbers of non-alveolar macrophages (includes interstitial macrophages and recruited macrophages) and DCs, which comprise the primary CCR2^+^ cells that traffic to the dLN to present Mtb antigen^33^, were lower in the lungs of *Irgm1*^-/-^ mice compared to WT mice at 21 dpi **(Fig. 1I)**. In contrast, the numbers of neutrophils and AMs were higher in the lungs of *Irgm1*^-/-^ mice compared to WT mice at 21 dpi **(Fig. 1J)**. The differences in cell populations in the lungs of *Irgm1*^-/-^ mice compared to WT mice were infection-dependent, as there were no significant differences in naïve uninfected mice **(Fig. S1)**. Analysis of Mtb burden in the different innate immune cell populations revealed a higher number of infected neutrophils and AMs, two cell types associated with poor control of Mtb replication^24,25,34,35^, in the lungs of *Irgm1^-/-^* mice compared to WT mice **(Fig. 1K-M)**. In addition, there was increased Mtb burden per neutrophil (based on Mtb-GFP mean fluorescence intensity (MFI)) at 21dpi in the lungs of *Irgm1^-/-^* mice compared to WT **(Fig. 1K, L)**. We were unable to reliably compare the Mtb burden in AMs because there were too few infected AMs in WT mice at 21 dpi. The increased numbers of Mtb-infected neutrophils and AMs at 21 dpi likely contribute to the higher Mtb burdens in the *Irgm1^-/-^* lungs at this time point.

To further evaluate inflammatory responses in *Irgm1^-/-^*mice during Mtb infection, we measured chemokine and cytokine levels in the lungs at 21 dpi using a cytokine bead array (Bio-Rad). The levels of IFN-γ, IL-12(p40), and RANTES were all lower at 21 dpi in the lungs of *Irgm1^-/-^* mice compared to WT mice **(Fig. 1N)**, consistent with the defect in T cell responses. In contrast, the lungs of Mtb-infected *Irgm1^-/-^* mice contained significantly higher levels of IL-6, KC, MCP-1, and MIP-1α **(Fig. 1O)**, all of which are associated with neutrophil inflammatory responses^36–39^. Together, the cytokine profiling data supports the observations of decreased activated CD4^+^ T cells and increased innate immune cell infection in the lungs of Mtb-infected *Irgm1^-/-^* mice.

### Adoptive transfer of activated Mtb-specific CD4^+^ T cell rescues control of Mtb infection in innate immune cells and allows for survival of *Irgm1*^-/-^ mice

The defect in T cell expansion in the dLN and resulting absence of activated T cells in the lungs of Mtb-infected *Irgm1^-/-^* mice was accompanied by increased numbers and infection of neutrophils and AMs in the lungs at 21 dpi. We, therefore, investigated whether restoring Mtb-specific T cell responses in *Irgm1*^-/-^ mice could rescue control of Mtb infection. To bypass the need for antigen presentation and T cell expansion *in vivo*, we adoptively transferred Mtb ESAT-6 specific CD4^+^ T cells that had been isolated from C7 TCR transgenic mice^40^ and activated *in vitro* by incubating with irradiated splenocytes, ESAT-6 peptide, α-IL-4, IL-12, and IL-2 **(Fig. 2A)**. *In vitro* activation of the C7 CD4^+^ T cells was confirmed by monitoring the expression of T-bet by flow cytometry **(Fig. 2B)**. We transferred 2.5 million activated C7 CD4^+^ T cells into each WT and *Irgm1^-/-^* mouse one day prior to Mtb infection and monitored immune cell infiltrate in the lungs at 21 dpi. The C7 cells express Thy1.1 and, thus, can be distinguished from the recipient mouse cells by flow cytometry **(Fig. S5B)**. *Irgm1^-/-^* mice accumulated a similar number of total C7 CD4^+^ T cells in the lung as WT mice at 21dpi **(Fig. 2C)**, demonstrating that *Irgm1^-/-^* mice were able to recruit activated Mtb-specific CD4^+^ T cells to the lung. Adoptive transfer of activated Mtb-specific C7 Thy1.1^+^ CD4^+^ T had no effect on the numbers of recipient-derived Thy1.1^-^ CD4^+^ T cells or activated Thy1.1^-^ CD4^+^ T cells in the lungs and dLN of *Irgm1*^-/-^ mice at 21 dpi **(Fig. S2A and S2B)**.

**Fig. 2:**
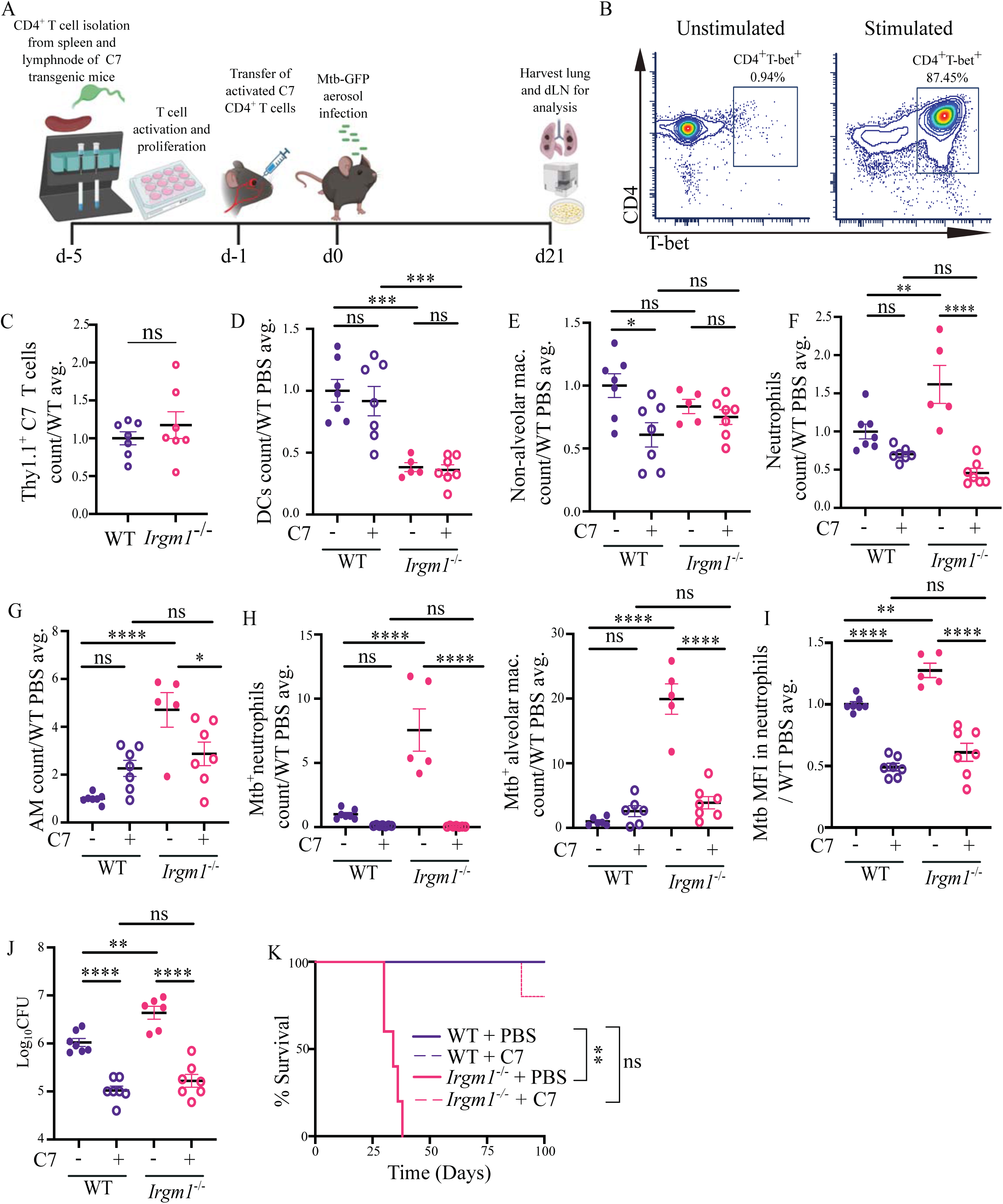
Adoptive transfer of activated Mtb-specific CD4^+^ T cell rescues control of Mtb infection in innate immune cells and allows for survival of *Irgm1*^-/-^ mice (A) Protocol schematic for activation and adoptive transfer of Mtb ESAT-6-specific C7 transgenic CD4^+^ T cells to mice prior to Mtb aerosol infection and analysis of immune cells at 21dpi. **(B)** Before transfer, isolated CD4^+^ T cells were analyzed for T-bet expression as a proxy of activation status after being stimulated *in vitro* with ESAT-6 peptide, α-IL-4, IL-12, and IL-2 or being left unstimulated. **(C-H)** Number of **(C)** C7 Thy1.1^+^CD4^+^ T cells, **(D)** DCs, **(E)** non-alveolar macrophages, **(F)** neutrophils, **(G)** AMs, and **(H)** Mtb^+^ neutrophils and AMs (based on GFP from Mtb-GFP) in the lungs of mice at 21 dpi that either received activated C7 CD4^+^ T cells (+) or PBS (-). **(I)** Mtb burden (based on MFI of GFP) per infected neutrophil in the lung at 21 dpi. **(J)** Mtb CFU count in lungs at 21 dpi. **(K)** Survival of mice following Mtb infection, where statistical significance was determined by Mantel-Cox test (**P=0.0017). (n=5 for each strain and condition). **(C-J)** Each data point corresponds to one mouse and graphs show the mean ± SEM, where in **(D-I)** the data are expressed as the ratio to the average for WT administered PBS in a single infection experiment to mitigate inter-experiment variation in total cell counts. Statistical significance of differences within a single mouse strain or treatment condition were determined by **(C)** two-tailed unpaired t-tests or **(D-J)** one-way ANOVA with Tukey’s multiple comparison test. **** P < 0.0001; *** P < 0.001; ** P < 0.01; * P < 0.05; ns=not significant.

The numbers of DCs remained low in the lungs of *Irgm1^-/-^*mice at 21 dpi following transfer of activated Mtb-specific CD4^+^ T cells **(Fig. 2D)** and although the number of non-alveolar macrophages were not significantly lower in *Irgm1^-/-^* mice compared to WT in these experiments, they remained unchanged in abundance following transfer of activated Mtb-specific CD4^+^ T cells **(Fig. 2E)**. These data suggest that T cell-independent processes contribute to the abundance of antigen-presenting cells in the lungs of *Irgm1^-/-^* mice. In contrast, transfer of activated Mtb-specific CD4^+^ T cells decreased the numbers of neutrophils and AMs in the lungs of *Irgm1^-/-^*mice at 21 dpi **(Fig. 2F, G)**, demonstrating that the presence of activated T cells is able to suppress the accumulation of these cell types in *Irgm1^-/-^* mice. Transfer of activated Mtb-specific CD4^+^ T cells also significantly reduced the number of Mtb-infected neutrophils and AMs in the lungs of *Irgm1^-/-^* mice at 21 dpi **(Fig. 2H)** as well as decreased the Mtb burden per neutrophil **(Fig. 2I)**. The improved control of Mtb infection in neutrophils and AMs following transfer of activated Mtb-specific CD4^+^ T cells corresponded to decreased Mtb burdens in the lungs of *Irgm1*^-/-^ mice at 21 dpi **(Fig. 2J)**. We also monitored the survival of WT and *Irgm1*^-/-^ mice that received activated Mtb-specific CD4^+^ T cells or PBS control and found that adoptive transfer of activated Mtb-specific CD4^+^ T cells significantly extended the survival in *Irgm1^-/-^* mice **(Fig. 2K)**, demonstrating that T cell-mediated control of Mtb infection in neutrophils and AMs is sufficient to rescue the susceptibility of *Irgm1*^-/-^ mice. Together these data indicate that IRGM1 expression is required for survival following Mtb infection in mice due to its role in supporting Mtb-specific T_H_1 responses that promote control of Mtb infection of AMs and neutrophils.

### Neutrophil depletion extends the survival of *Irgm1^-/-^* mice during Mtb infection

The extension in survival of Mtb-infected *Irgm1*^-/-^ mice following adoptive transfer of activated Mtb-specific CD4^+^ T cells was associated with decreased numbers of Mtb-infected neutrophils in the lungs at 21 dpi **(Fig. 2H, I)**. Excessive neutrophil recruitment and infection is associated with Mtb replication, tissue damage, and progression of Mtb disease in mice and humans^16,41–47^. If the critical role for T_H_1 responses that is lost in *Irgm1*^-/-^ mice during Mtb infection is control of Mtb infection and replication in neutrophils, then depletion of neutrophils might prolong survival of Mtb-infected *Irgm1*^-/-^ mice. To directly test this possibility, we administered a neutrophil-depleting αLy6G (clone 1A8) antibody or isotype control antibody to WT and *Irgm1*^-/-^ mice beginning on 10 dpi with repeated injections every other day until 34 dpi **(Fig. S3A)** and monitored survival over time. Transiently depleting neutrophils in Mtb-infected *Irgm1^-/-^* mice from 10-34 dpi extended their median survival time from 43 dpi for IgG-treated *Irgm1*^-/-^ mice to 71 dpi for α-Ly6G-treated *Irgm1*^-/-^ mice **(Fig. 3A)**, indicating that neutrophils directly contribute to the susceptibility of *Irgm1^-/-^*mice to Mtb.

**Fig. 3:**
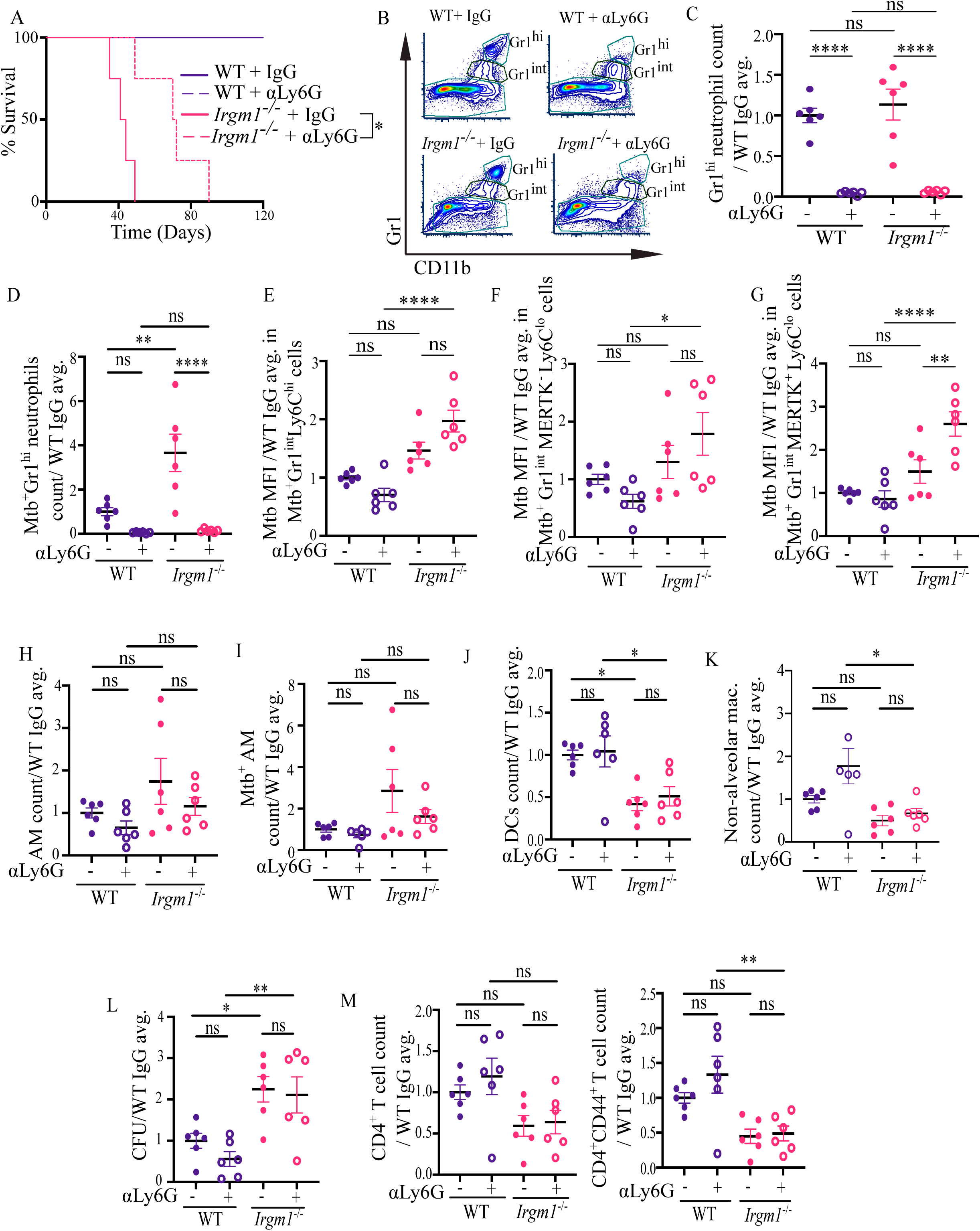
Neutrophil depletion extends the survival of *Irgm1^-/-^* mice during Mtb infection. **(A)** Survival of WT and *Irgm1^-/-^*mice following Mtb aerosol infection and administration of an αLy6G antibody to deplete neutrophils or an IgG control antibody (n=4 for each strain and condition). Statistical significance of difference in survival between *Irgm1^-/-^*mice treated with αLy6G antibody versus IgG control was determined by Mantel-Cox test (*P<0.05). **(B)** Representative flow plot showing successful depletion of Gr1^hi^ neutrophils, but not Gr1^int^ cells, upon αLy6G administration in both WT and *Irgm1^-/-^* mice. **(C-D)** Number of **(C)** Gr1^hi^ neutrophils and **(D)** Mtb^+^ Gr1^hi^ neutrophils in the lungs at 21dpi following administration of αLy6G antibody (+) or an IgG control antibody (-). **(E-G)** Mtb burden (based on MFI of GFP) per infected cells in **(E)** Mtb^+^Gr1^int^Ly6C^hi^ **(F)** Mtb^+^Gr1^int^MERTK^-^Ly6C^lo^ and **(G)** Mtb^+^Gr1^int^MERTK^+^Ly6C^lo^ cells at 21dpi in the WT and *Irgm1*^-/-^ mouse lung following administration of αLy6G antibody (+) or IgG (-). **(H-K)** Number of **(H)** AMs, **(I)** Mtb^+^ AMs, **(J)** DCs, and **(K)** non-alveolar macrophages in the lungs at 21dpi. **(L)** Mtb CFU count in the lung at 21dpi per WT IgG average; CFU values were presented as a ratio to mitigate the variation between experiments and the raw values are shown in Fig. S3I. **(M)** Number of CD4^+^ T cells and activated CD4^+^ T cells in the lungs at 21 dpi. Each data point corresponds to one mouse and graphs show the mean ± SEM, where in **(C-M)** the data are expressed as the ratio to the average for WT administered IgG control antibody in a single infection experiment to mitigate inter-experiment variation in total cell counts. **(C-M)** Statistical significance of differences within a single mouse strain or treatment condition were determined by one-way ANOVA’s Tukey’s multiple comparison test. **** P < 0.0001; ** P < 0.01; * P < 0.05. ns=not significant.

We monitored the effects of neutrophil depletion on immune cell infiltrate and Mtb burden at 21 dpi to identify the changes associated with the extended survival. We confirmed the efficient depletion of Gr1^hi^ neutrophils (Ly6C^lo^) with αLy6G antibody treatment in the lungs of WT and *Irgm1*^-/-^ mice at 21 dpi **(Fig. 3B, C)**, however, we also noted the persistence of a Gr1 intermediate (Gr1^int^) population that was resistant to depletion with the αLy6G antibody **(Fig. 3B)**. The Gr1^int^ population had two distinct populations of cells that either expressed high or low levels of Ly6C (Ly6C^hi^ or Ly6C^lo^) on the cell surface **(Fig. S3B)** where the Ly6C^hi^ cells are likely inflammatory monocytes and the Ly6C^lo^ cells are likely non-classical monocytes or granulocytes, based on prior publications^48–50^. To separate the Ly6C^lo^ non-classical monocyte and granulocyte populations, we gated the Ly6C^lo^ cells based on surface expression of the monocyte/macrophage marker MERTK^51^ **(Fig. S3B)**. Administration of αLy6G antibody decreased the number of MERTK^+^ Ly6C^lo^ GR1^int^ cells in the lungs of both WT and *Irgm1*^-/-^ mice compared to the IgG controls, whereas, the Ly6C^hi^ Gr1^int^ and MERTK^-^ Ly6C^lo^ Gr1^int^ cells remained unchanged following αLy6G treatment **(Fig. S3C-S3E)**. There were no significant differences in the numbers of any of the Gr1^int^ cell populations between WT and *Irgm1*^-/-^ mice regardless of IgG or αLy6G treatment **(Fig. S3C-E)**. Administration of αLy6G antibody decreased the number of Mtb infected Gr1^hi^ neutrophils in the lungs of *Irgm1*^-/-^ mice **(Fig. 3D)** but did not change the numbers of Mtb-infected Gr1^int^ cells **(Fig. S3F-H)**. However, administration of αLy6G antibody resulted in the Mtb burden per cell for each Gr1^int^ population being significantly higher in *Irgm1*^-/-^ mice compared to WT **(Fig. 3E-G)**. αLy6G antibody treatment in Mtb-infected *Irgm1*^-/-^ mice also resulted in a trend of lower numbers of AMs and infected AMs, however, none of the differences for this population were statistically significant in these experiments **(Fig. 3H, I)**. αLy6G treatment did not rescue the lower numbers of DCs or non-alveolar macrophages in *Irgm1^-/-^*mice at 21 dpi **(Fig. 3J, K)**, indicating that these lower numbers of these cell types occurs independently of the higher numbers of Gr1^hi^ neutrophils. Despite the increased survival of Mtb-infected *Irgm1*^-/-^ mice following Gr1^hi^ neutrophil depletion, the lung Mtb burdens remained high **(Fig. 3L and Fig. S3I)**, which was likely due to the maintained high levels of Mtb infection in the Gr1^int^ cells.

Since transfer of activated Mtb-specific CD4^+^ T cells was sufficient to promote control of Mtb infection in neutrophils, we analyzed CD4^+^ T cell numbers at 21 dpi following αLy6G antibody treatment to determine if the T cell abundance remained low in *Irgm1*^-/-^ mice and this accounted for the high levels of Mtb infection in Gr1^int^ cells. Indeed, αLy6G antibody treatment did not alter CD4^+^ T cell numbers or activation in the lungs of *Irgm1^-/-^* mice at 21 dpi **(Fig. 3M)**, likely accounting for the continued high level of Gr1^int^ cell infection in αLy6G treated *Irgm1^-/-^* mice. Together these data indicate that neutrophils contribute to the susceptibility of *Irgm1^-/-^* mice to Mtb infection, and their contribution is downstream of defects in CD4^+^ T cell expansion in Mtb-infected *Irgm1*^-/-^ mice.

### Increased type I IFN signaling in *Irgm1^-/-^*mice causes defects in T cell expansion and susceptibility during Mtb infection

To investigate what pathways are regulated by IRGM1 that could be contributing to control of Mtb pathogenesis, we performed bulk RNA sequencing (RNAseq) analysis on total lung from WT and *Irgm1*^-/-^ mice at 21dpi. There were 2372 differentially expressed genes (p_adj._ :≤ 0.05), out of which 1072 were downregulated **(Table S1)** and 1300 were upregulated **(Table S2)**, in the lungs of *Irgm1*^-/-^ mice compared to WT at 21 dpi. We determined the pathways enriched for genes that were significantly upregulated in the absence of *Irgm1* by inputting the list of significantly upregulated genes into EnrichR and querying the GO Biological Process-2021 gene set database^52,53^. The top 10 pathways that were significantly positively enriched in *Irgm1*^-/-^ mice at 21 dpi were all related to IFN signaling and the top 7 out of 10 pathways were specifically related to type I IFN **(Fig. 4A)**. In addition, the genes within the “Cellular Response to Type I Interferon” GO pathway (GO:0071357) were amongst the highest upregulated genes in the lungs of *Irgm1*^-/-^ mice at 21 dpi **(Fig. S4A)**. We confirmed the upregulation of *Ifnb1* (gene for IFN-β) and several IFN stimulated genes (ISGs) in the lungs of *Irgm1*^-/-^ mice at 21 dpi by quantitative real-time PCR (qRT-PCR) **(Fig. 4B)**. Previous reports have shown that the lungs of naïve *Irgm1^-/-^* mice exhibit elevated expression of type I IFNs and ISGs and this improved host survival during viral infections^54–56^. We also observed increased ISG expression in the lungs of naïve *Irgm1*^-/-^ mice compared to WT controls by RNAseq and qRT-PCR, supporting that type I IFN signaling was elevated in *Irgm1*^-/-^ mice even before Mtb infection **(Table S3 and Fig. S4B)**. Type I IFN has been associated with poor control of Mtb infection in humans and mice^47,57–60^, although the mechanistic basis for how type I IFN signaling interferes with control of Mtb remains unknown. In addition, deletion of the gene encoding the type I IFN receptor, *Ifnar1*, in *Irgm1*^-/-^ mice rescues their susceptibility to Mtb infection through 150 dpi **(Fig. 4C)**^61^, indicating that increased type I IFN signaling contributes to the susceptibility of *Irgm1*^-/-^ mice to Mtb.

**Fig. 4:**
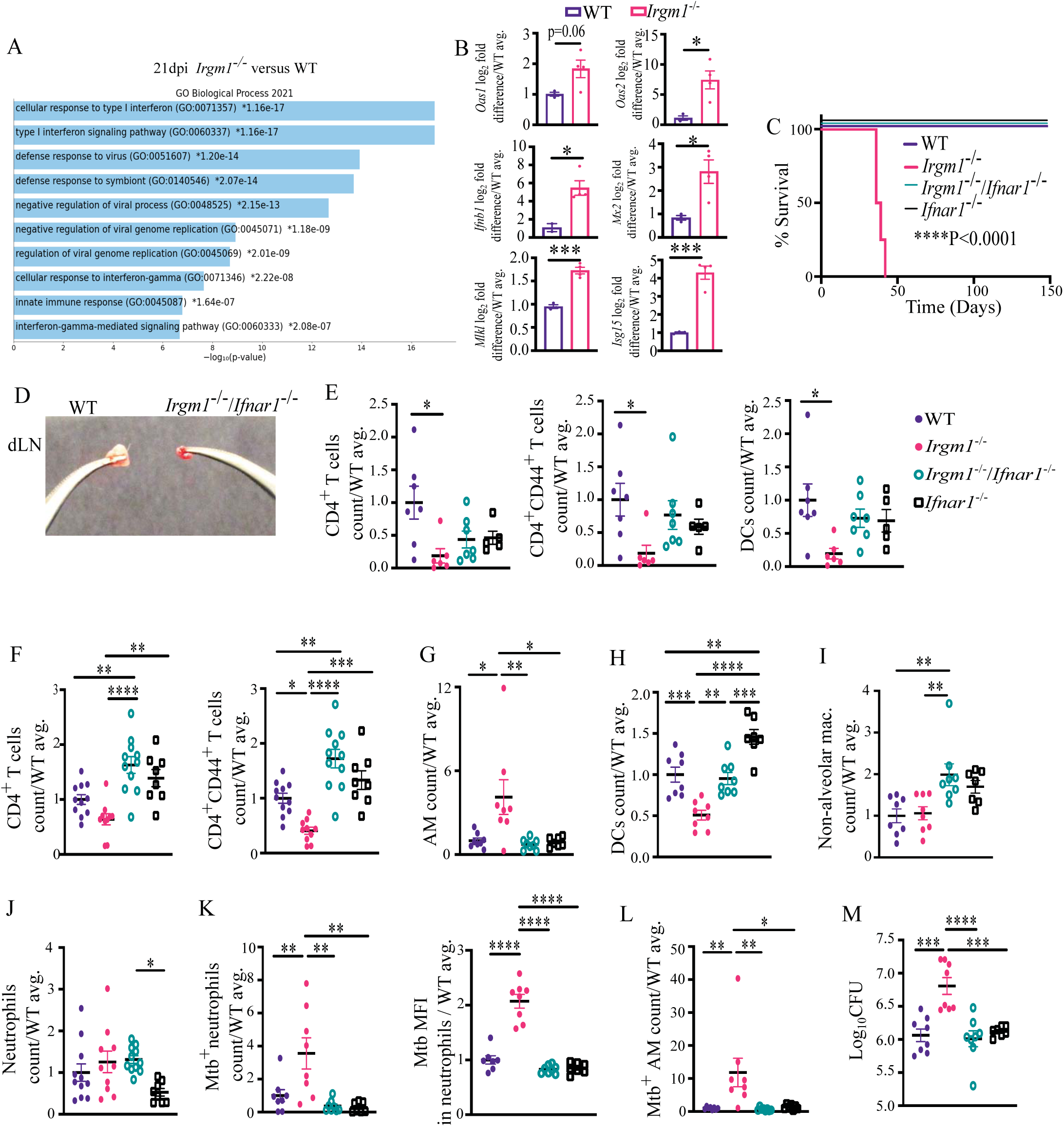
Increased type I IFN signaling in *Irgm1^-/-^* mice causes defects in T cell expansion and susceptibility during Mtb infection. **(A)** Ten most enriched pathways in the upregulated genes from bulk RNAseq analysis of total lung from *Irgm1^-/-^* mice compared to WT controls at 21 dpi. **(B)** qRT-PCR analysis of ISG expression in lungs of *Irgm1^-/-^*mice compared to WT controls at 21 dpi. Significance of differences were determined by two-tailed unpaired t-test. *** P < 0.001; * P < 0.05. **(C)** Survival of mice following Mtb infection (WT n=5, *Irgm1*^-/-^ n=4, *Irgm1*^-/-^/*Ifnar1*^-/-^ n=9, *Ifnar1*^-/-^ n=5). Statistical significance of differences in survival compared to *Irgm1^-/-^* mice determined by Mantel-Cox test. **(D)** Representative mediastinal dLN from WT and *Irgm1^-/-^*/*Ifnar1*^-/-^ mice at 21 dpi. **(E)** Number of CD4^+^ T cells, activated CD4^+^ T cells (CD44^+^), and DCs in the dLN at 21dpi. The legend in **(E)** applies to all subsequent panels. **(F-L)** Number of **(F)** CD4**^+^** T cells and activated CD4^+^ (CD44^+^CD62L^-^) T cells, **(G)** AMs, **(H)** DCs, **(I)** Non-alveolar macrophages, **(J)** neutrophils, **(K)** Mtb^+^ neutrophils and Mtb burden (based on MFI of GFP) in Mtb-infected neutrophils, and **(L)** Mtb^+^ AMs in the lung at 21dpi. **(M)** Mtb CFU count in the lung at 21dpi. Each data point corresponds to one mouse and graphs show the mean ± SEM, where in **(E-L)** the data are expressed as the ratio to the average for WT mice. Statistical significance of differences was determined by one-way ANOVA’s Tukey’s multiple comparison test and only significant comparisons are shown. **** P < 0.0001; *** P < 0.001; ** P < 0.01; * P < 0.05.

To identify correlates of protection when type I IFN signaling is blocked in *Irgm1*^-/-^ mice, we compared the immune cell composition and Mtb burdens in WT, *Irgm1*^-/-^, *Irgm1*^-/-^*/Ifnar1*^-/-^, and *Ifnar1*^-/-^ mice at 21 dpi. When we analyzed the dLN at 21 dpi, we found that, despite the complete rescue in survival, the *Irgm1*^-/-^*/Ifnar1*^-/-^ mice still had smaller dLN than WT mice at 21 dpi **(Fig. 4D)**, where ablating type I IFN signaling only intermediately increased the numbers of CD4^+^ T cells, activated CD4^+^ T cells, and DCs in the dLN **(Fig. 4E)**. However, ablating type I IFN signaling increased the abundance of CD4^+^ T cells and activated CD4^+^ T cells in the lungs of *Irgm1*^-/-^ mice to numbers even higher than in WT mice **(Fig. 4F)**. The restored T cell responses in the lungs of *Irgm1*^-/-^ mice when *Ifnar1* was deleted was associated with decreased AMs **(Fig. 4G)**, increased DCs **(Fig. 4H),** and increased nonalveolar macs **(Fig. 4I).** Deletion of *Ifnar1* in *Irgm1*^-/-^ mice also resulted in lower numbers of Mtb-infected neutrophils and lower burdens of Mtb per neutrophil, without affecting the total number of neutrophils in the lungs **(Fig. 4J, K)**. Blocking type I IFN signaling in *Irgm1*^-/-^ mice similarly led to fewer Mtb infected AMs in the lung **(Fig. 4L)**. The decreased infection of neutrophils and AMs correlated with decreased Mtb burden in the lungs of *Irgm1*^-/-^*/Ifnar1*^-/-^ mice compared to the *Irgm1*^-/-^ mice **(Fig. 4M)**. Therefore, loss of type I IFN signaling reversed most phenotypes in the lungs of Mtb-infected *Irgm1*^-/-^ mice.

These studies indicate that the defective lung T_H_1 responses and increased neutrophil infection are downstream of the enhanced type I IFN signaling in Mtb-infected *Irgm1*^-/-^ mice. To directly test this, we analyzed ISG expression in the lungs of WT and *Irgm1*^-/-^ mice at 21 dpi following transfer of activated Mtb-specific CD4^+^ T cells or neutrophil depletion **(Fig. 2A and Fig. S3A)**. Despite the improved survival following transfer of activated Mtb-specific CD4^+^ T cells **(Fig. 2K)** or neutrophil depletion **(Fig. 3A)**, ISG expression remained high in the lungs of Mtb-infected *Irgm1*^-/-^ mice **(Fig. S4C, D)**. Therefore, loss of T cell expansion and increased neutrophil infection were consequences, but not drivers, of the higher levels of type I IFN in *Irgm1*^-/-^ mice during Mtb infection.

### Increased type I IFN signaling in *Irgm1*^-/-^ mice during Mtb infection is dependent on IRGM3 expression

Deletion of *Irgm3* in *Irgm1*^-/-^ mice rescues the higher bacterial burdens and early susceptibility during Mtb infection^5^ **(Fig. 5A, B)**, but the mechanistic basis of the rescue is unknown. IRGM1 has been shown to maintain mitochondrial quality control and suppress mitochondrial DNA-induced type I IFN production in the presence of IRGM3 expression in fibroblasts and macrophages^62,63^. As such, deletion of *Irgm1* in macrophages and fibroblasts leads to increased production of type I IFN, which is reversed if *Irgm3* is also deleted^62^. Therefore, we investigated whether the increased type I IFN signaling observed in *Irgm1*^-/-^ mice during Mtb infection was dependent on IRGM3 expression by infecting *Irgm1*^-/-^/*Irgm3*^-/-^ mice and measuring ISG expression in the lungs at 21 dpi. We observed a significant decrease in the expression of ISGs in the lungs of *Irgm1*^-/-^/*Irgm3*^-/-^ mice compared to the *Irgm1*^-/-^ mice **(Fig. 5C)**, indicating that indeed the increased type I IFN in *Irgm1*^-/-^ mice is dependent on IRGM3 expression. Similar to as observed when *Ifnar1* was deleted, the decreased type I IFN signaling in the *Irgm1*^-/-^/*Irgm3*^-/-^ mice was associated with increased numbers of activated CD4^+^ activated T cells in the dLN **(Fig. 5D)** and lungs **(Fig. 5E)**, increased DC numbers in the dLN **(Fig. 5F)** and lung **(Fig. 5G)**, and decreased infection of neutrophils **(Fig. 5H)** and AMs **(Fig. 5I)**. Therefore, the data suggest that IRGM3-induced type I interferon-mediated immunodeficiency in the absence of *Irgm1* results in susceptibility to Mtb infection in mice.

**Fig. 5:**
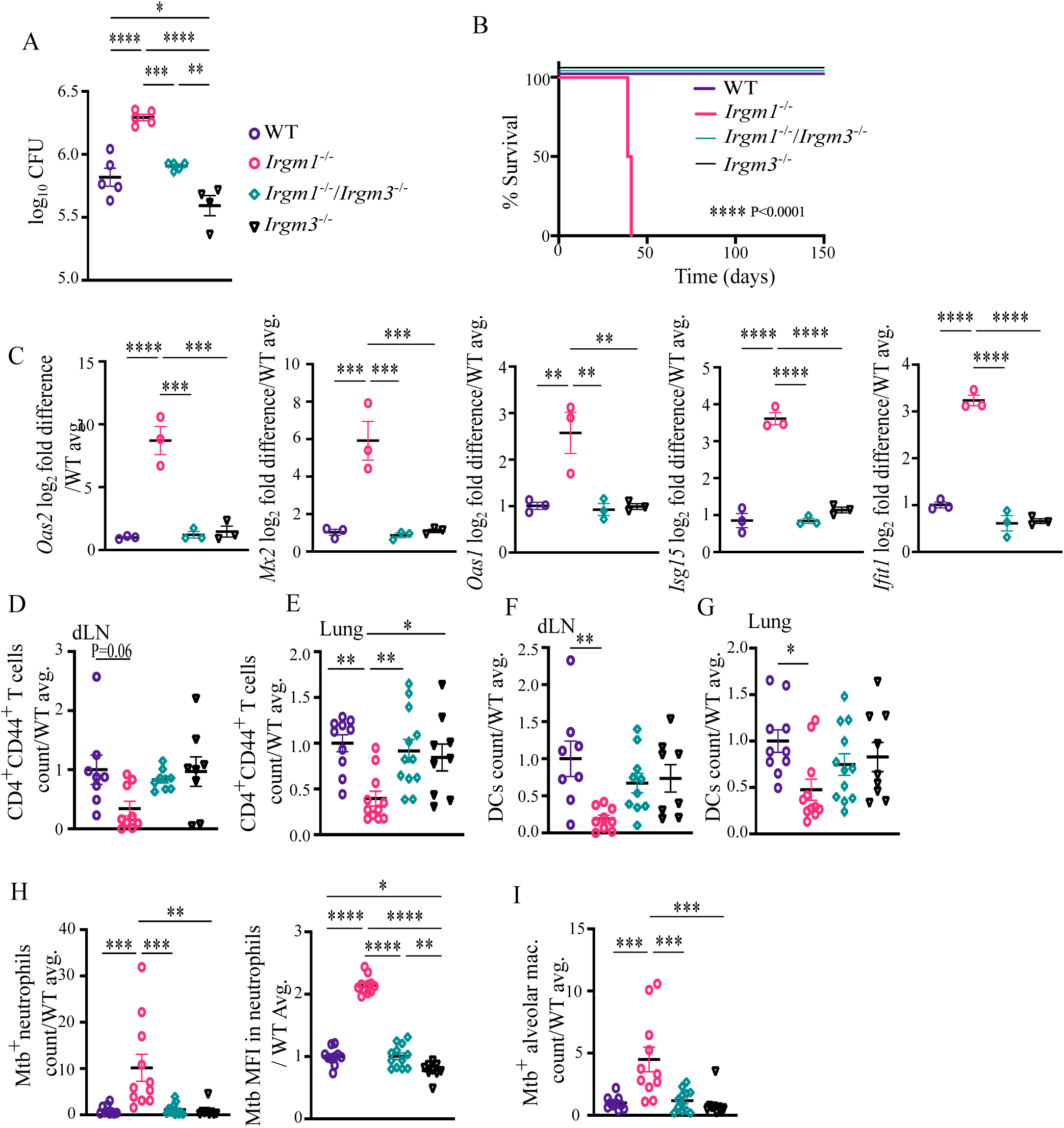
Increased type I IFN signaling in *Irgm1*^-/-^ mice during Mtb infection is dependent on IRGM3 expression. **(A)** Mtb CFU count in lungs at 21 dpi. The legend in **(A)** applies to all subsequent panels. **(B)** Survival of mice following Mtb infection (WT n=3, *Irgm1*^-/-^ n=2, *Irgm1*^-/-^/*Irgm3*^-/-^ n=8 and *Irgm3*^-/-^ n=8). Statistical significance of differences in survival compared to *Irgm1^-/-^* mice determined by Mantel-Cox test. **(C)** qRT-PCR analysis of ISG expression in lungs at 21 dpi. **(D-E)** Number of activated CD4^+^ T cells (CD44^+^) in **(D)** dLN and **(E)** lungs at 21dpi. **(F-G)** Number of DCs in **(F)** dLN and **(G)** lungs at 21dpi. **(H-I)** Number of **(H)** Mtb^+^ neutrophils and Mtb burden (based on MFI of GFP) in Mtb infected neutrophils and **(I)** Mtb^+^ AMs in the lung at 21dpi. Each data point corresponds to one mouse and graphs show the mean ± SEM, where in **(D-I)** the data are expressed as the ratio to the average for WT mice. Statistical significance of differences was determined by one-way ANOVA’s Tukey’s multiple comparison test and only significant comparisons are shown. **** P < 0.0001; *** P < 0.001; ** P < 0.01; * P < 0.05.

## DISCUSSION

In this manuscript, we uncover the biological basis for how IRGM1 promotes control of Mtb infection. A role for IRGM1 in a protective host response to Mtb infection was identified over two decades ago^6^ and initially the focus was on IRGM1 facilitating targeting of Mtb to lysosomes through the process of autophagy^6,12–14^. However, this model was drawn into question following reports that IRGM1 did not colocalize with mycobacteria in IFN-γ-treated cells^15^ and the discovery that autophagy was not required to control Mtb replication in macrophages during infection of mice *in vivo*^16,17^. More recent work has associated the requirement for IRGM1 during Mtb infection with IRGM3 activity^5^, type I IFN signaling^61^, and T_H_1 responses^18^, however, it remained unknown how these pathways intersected to impact IRGM1-mediated control of Mtb infection. Based on our data, we propose a model **(Fig. 6)** wherein IRGM1 is required to suppress IRGM3-mediated induction of type I IFN in naïve mice. The higher levels of type I IFN in the absence of IRGM1 condition the environment of the mouse so that following infection, Mtb-specific T cells fail to expand and antigen presenting cells, such as DCs, decrease in numbers. The absence of T_H_1 responses allows for uncontrolled Mtb infection and replication in neutrophils and AMs, which directly contributes to the increased susceptibility of *Irgm1*^-/-^ mice to Mtb infection. These studies identify the mechanistic link between IRGM1, IRGM3, type I IFN, and T_H_1 responses during Mtb infection, while also revealing new roles for type I IFN signaling in suppressing the DC accumulation and T_H_1 responses in the lungs that are critical for promoting control of Mtb replication in neutrophils.

**Fig. 6:**
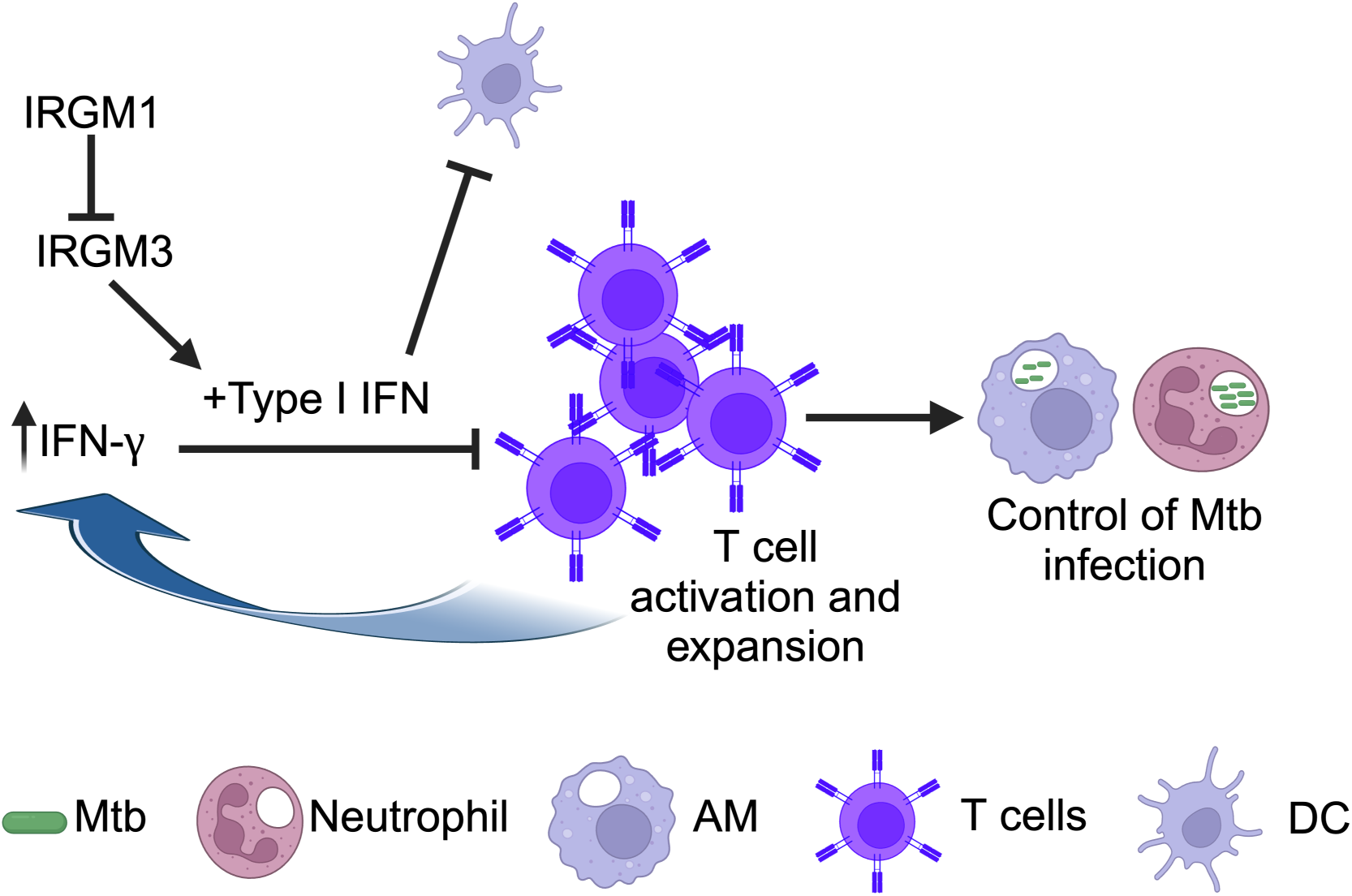
Model of IRGM1-mediated control of Mtb pathogenesis. Schematic diagram of model where IRGM1 is required to suppress IRGM3-mediated induction of type I in naïve mice. Following Mtb infection, the higher levels of type I IFN sensitize activated T cells to IFN-γ-stimulated cell death, precluding T_H_1 CD4^+^ T cell expansion. In addition, the heightened type I IFN signaling leads to decreased numbers of DCs in the lungs and dLN during Mtb infection, which may also negatively impact the initiation of T_H_1 responses. Loss of T_H_1 CD4^+^T cell expansion results in uncontrolled Mtb infection and replication in neutrophils and AMs, causing the mice to succumb to the infection.

Type I IFN is associated with poor control of Mtb infection in humans and mice^43,47,58–60^, however, it remains unknown the mechanisms by which type I IFN could be suppressing control of Mtb. Given that the susceptibility of *Irgm1*^-/-^ mice to Mtb infection is dependent on type I IFN signaling, better understanding of the pathways driving the susceptibility of these mice could shed light on how more generally type I IFN impacts protective immune responses to Mtb. In addition, there are multiple features of the *Irgm1*^-/-^ mice that bear similarity to STING gain-of-function mutants, which also exhibit constitutively high levels of type I signaling^64–66^. Mice with STING gain-of-function mutations develop T cell-dependent lung disease^67^ and also lack lymph nodes, which is attributed to decreased numbers of innate lymphoid cells, including lymphoid tissue inducer cells, in STING gain-of-function mice^68^. Although, it is not known why there are defects in the numbers of lymphoid tissue inducer cells in STING gain-of-function mice, it is possible that the mechanism is shared with *Irgm1*^-/-^ mice and may be related to heightened type I IFN signaling in both of these mouse backgrounds. In addition, IFN-γ signaling is required for the lymph node development and T cell defects in STING gain-of-function mice^69^, similar to our model that elevated type I IFN signaling in *Irgm1*^-/-^ mice sensitizes the CD4^+^ T cells to IFN-γ-dependent cell death, where IFN-γ is produced upon activation during Mtb infection. One could also imagine that the type I IFN conditioning of the tissue that occurs in naïve *Irgm1*^-/-^ mice could represent a similar scenario as when a viral infection proceeds Mtb infection, as could happen in HIV-infected individuals who are exposed to Mtb. Whether there are aspects of *Irgm1*^-/-^ mice that model type I IFN induction due to HIV, or other viral, infections is an area for future investigations.

The studies herein used germline deletion of *Irgm1* and it still remains unknown what cell types require *Irgm1* expression for control of Mtb infection. *Irgm1*-deficient T cells were able to rescue the susceptibility of *RAG2*^-/-^ mice to *M. avium* infection, suggesting that the defect in survival following activation was not T cell intrinsic^18^. However, *in vitro* studies revealed that IRGM1-deficient T cells exhibit decreased survival and defective expansion following activation in the presence of IFN-γ, suggesting that there are T cell intrinsic roles for IRGM1 as well as roles for IRGM1 in other cells that contribute to T cell survival following activation in the presence of IFN-γ^19^. Cell type specific effects of IRGM1 expression on type I IFN signaling have also been described, where deletion of *Irgm1* in fibroblasts leads to increased type I IFN due to a defect in mitophagy that is associated with mitochondrial DNA-induced type I IFN responses via the cGAS-STING pathway^62^. In contrast, deletion of *Irgm1* in macrophages leads to increased type I IFN signaling via lysosomal toll-like receptors^62^. In addition, human IRGM promotes autophagy-mediated degradation of nucleic acid sensors to restrain type I IFN responses in macrophages and epithelial cells^63^. Therefore, it is likely that IRGM1 has cell-type specific effects that could differentially impact control of Mtb infection.

In humans, a polymorphism in the *IRGM* gene that decreases its expression has been associated with active TB disease in humans^3,4^ and given the enhanced susceptibility *Irgm1*^-/-^ mice to Mtb infection, it is possible there are parallels between the roles for IRGM1 in mice with IRGM in humans. However, interpretating the relationship between human and mouse IRGM homologs is complicated by the expansion of the IRGM family in mice. In mice, IRGM3 often plays a key role in the effects of *Irgm1* deletion. In addition to its role in autophagy^10,11^, IRGM3 regulates endoplasmic reticulum (ER) stress responses in a PI3K/AKT-dependent manner during Coxsackievirus B3 infection^70^, where ER stress responses are associated with the induction of type I IFN signaling^71,72^. The contribution of ER stress responses and other yet to be identified roles for IRGM3 in host responses to Mtb infection remains an open question. Understanding the network of IRGM homologs will shed light on cell stress responses and type I IFN signaling during infection.

## MATERIALS AND METHODS

### Mice

WT, *Irgm1^-/-^*, *Ifnar1^-/-^*, *Irgm1^-/-^Ifnar1^-/-^*, *Irgm3^-/-^*, *Irgm1^-/-^Irgm3^-/-^* mice were housed and bred at Washington University in St. Louis specific pathogen-free conditions in accordance with federal and university guidelines, and protocols were approved by the Animal Studies Committee of Washington University. *Irgm1^-/-^* and *Ifnar1^-/-^* mice were a gift from Dr. Ashley Steed^54,73^ at Washington University in St. Louis. *Irgm3*^-/-^ mice were purchased from Jackson Laboratory (B6.129-*Igtp^tm1Gvw^*/J; 034702). C7 TCR tg/thy1.1 mice were a gift from Dr. Michael Glickman at Sloan Kettering Institute^40^. All the mice used for experimental procedures were backcrossed to C57BL/6. Male and female littermates (aged 8–12 weeks) were used and were subject to randomization. All procedures involving animals were conducted following the National Institutes of Health (NIH) guidelines for housing and care of laboratory animals and performed in accordance with institutional regulations after protocol review and approval by the Institutional Animal Care and Use Committee of the Washington University in St. Louis School of Medicine. Washington University is registered as a research facility with the United States Department of Agriculture and is fully accredited by the American Association of Accreditation of Laboratory Animal Care. The Animal Welfare Assurance is on file with Office for Protection from Research Risks–NIH. All animals used in these experiments were subjected to no or minimal discomfort. All mice were euthanized by CO_2_ asphyxiation, which is approved by the American Veterinary Association Panel on Euthanasia.

### Mtb infection in mice and determination of lung burden

Mtb Erdman or GFP-Mtb^31,32^ Erdman was cultured at 37° C in 7H9 (broth) or 7H11 (agar) (Difco) medium supplemented with 10% oleic acid/albumin/dextrose/catalase (OADC), 0.5% glycerol, and 0.05% Tween 80 (broth). GFP-Mtb was grown in the presence of kanamycin to ensure plasmid retention. Mtb cultures in logarithmic growth phase (OD600 = 0.5-0.8) were washed with PBS + 0.05% Tween-80, sonicated to disperse clumps, and diluted in sterile water. Mice were infected with 100-200 CFUs of aerosolized Mtb per mouse using an Inhalation Exposure System (Glas-Col). Within 2h of each infection, lungs were harvested from at least two control mice, homogenized, and plated on 7H11 agar to determine the input CFU dose. At each time point after infection, Mtb titers were determined by homogenizing the right lung’s superior, middle, and inferior lobes and plating serial dilutions on 7H11 agar. After 3 weeks of incubation at 37°C in 5% CO2, colonies were counted.

### Flow cytometry

Lungs were perfused with sterile PBS and the left lobes of the lungs were digested at 37°C with 630 µg/ml collagenase D (Roche) and 75 U/ml DNase I (Sigma). Single-cell suspensions were stained for live-dead Zombie NIR dye (BioLegend) (1:2000) followed by preincubation with Fc Block antibody (BD PharMingen) in PBS + 2% heat-inactivated FBS for 10 min at room temperature before staining. Cells were incubated with the following antibodies: Tcr-β-BV421 (BioLegend #109229;clone H57-597), CD19-pacific blue (BioLegend#115523;clone 6D5), CD44-BV510 (BioLegend#103044;clone IM7), CD4-BV570 (BioLegend#100541;clone RM4-5), Ly-6C-BV605 (BioLegend#128036;clone HK1.4), CD11b-BV650 (BioLegend#101259;clone M1/70), Ly-6G-BV785 (BioLegend#127645;clone 1A8), MHCII-Sparkblue (BioLegend#107662; clone M5/114.15.2), CD11c-PerCP (BioLegend#117325;clone N418), CD64-PerCPefluor710 (Invitrogen#46-0641-82;clone X54-5/7.1), SiglecF-PE (BD#552126;clone E50-2440), CD62L-PE-Cy5 (BioLegend#104410;clone MEL-14), MERTK-PE-Cy7 (Invitrogen#25-5751-82;clone DS5MMER), CD45-AF700 (BioLegend#103128;clone 30-F11), CD8a-APCfire750 (BioLegend#100766;clone 53-6.7). Cells were stained for 20 min at 4°C, washed, and fixed in 4% paraformaldehyde in PBS for 20 min at room temperature. After incubation, cells were washed again with FACS buffer and analyzed on a Cytek Aurora Spectral Flow Cytometer with 4L16V-14B-10YG-*r configuration using SpectroFlo V2.2.0.3 (Cytek). Absolute cell counts were determined by volumetric-based counting on the Cytek Aurora. Flow cytometry data were analyzed using FCS Express 7.12.0005. Gating strategies are depicted in **Fig. S5**.

### Cytokine analysis

For cytokine analysis, Lungs were homogenized in 5 ml of PBS supplemented with 0.05% Tween 80. Homogenates were filtered (0.22 µm) twice and analyzed by BioPlex-Pro Mouse Cytokine 23-Plex Immunoassay (Bio-Rad). Lung homogenates were quantified for IL-1α, IL-β, IL-2, IL-3, IL-4, IL-5, IL-6, IL-9, IL-10, IL-12p40, IL-13, IL-17A, Eotaxin, G-CSF, GM-CSF, IFN-γ, KC, MCP-1, MIP-1α, MIP-1β, RANTES, TNF-α. Cytokines not shown in Figure 1 were not significantly different between WT and *Irgm1^-/-^* mice or were out of the detection range.

### Activated T cell transfer and neutrophil depletion

ESAT6-specific C7 T cells were isolated from lymph nodes and spleen from C7 TCR tg/thy1.1 transgenic mice with Miltenyi Biotec CD4^+^ T cell isolation kit (130-104-454) and the cell purity (95-97%) was assessed by flow cytometry. C7 T cells were activated *in vitro* with ESAT-6 (5μg/ml), anti-IL4 (BioXcell #BE0045 clone 11B11;5μg/ml), and rIL12 (20ng/ml) for two days followed by co-culturing with 50U/ml IL-2 for an additional two days. After four days, T cell activation was assessed by flow cytometry for the expression of T-bet. 2.5 million activated T cells in PBS or the same volume of PBS were transferred intravenously to each mouse one day prior to aerosol infection with Mtb. For neutrophil depletion, mice were intraperitoneally injected every 48 h with 200 µg monoclonal anti-Ly6G antibody (clone 1A8; Leinco) or 200 µg polyclonal rat serum IgG (Sigma-Aldrich) diluted in sterile PBS beginning at 10 dpi and ending at 34 dpi.

### RNA isolation, RNAseq, and qRT-PCR analysis

RNA was extracted by lysing bulk lung tissue in TRIzol reagent (Invitrogen) using zirconia/silica beads with a FastPrep-24 bead beater (MP Biomedicals) followed by extraction using chloroform. RNA was isolated as per the manufacturer’s instruction using Direct-zol RNA Miniprep Plus kit (Zymo Research).

For RNAseq, RNA samples were prepared according to library kit manufacturer’s protocol, indexed, pooled, and sequenced on an Illumina NovaSeq 6000. Basecalls and demultiplexing were performed with Illumina’s bcl2fastq software and a custom python demultiplexing program with a maximum of one mismatch in the indexing read. RNA-seq reads were then aligned to the Ensembl release 76 primary assembly with STAR version 2.5.1a^74^. Gene counts were derived from the number of uniquely aligned unambiguous reads by Subread:featureCount version 1.4.6-p5^75^. Isoform expression of known Ensembl transcripts were estimated with Salmon version 0.8.2. Sequencing performance was assessed for the total number of aligned reads, total number of uniquely aligned reads, and features detected. The ribosomal fraction, known junction saturation, and read distribution over known gene models were quantified with RSeQC version 2.6.2^76^. All gene counts were then imported into the R/Bioconductor package EdgeR^77^ and TMM normalization size factors were calculated to adjust for samples for differences in library size. The TMM size factors and the matrix of counts were then imported into the R/Bioconductor package Limma^78^. Weighted likelihoods based on the observed mean-variance relationship of every gene and sample were then calculated for all samples with Limma’s voomWithQualityWeights^79^. The performance of all genes was assessed with plots of the residual standard deviation of every gene to their average log-count with a robustly fitted trend line of the residuals. Differential expression analysis was then performed to analyze for differences between conditions and the results were filtered for only those genes with Benjamini-Hochberg false-discovery rate adjusted p-values less than or equal to 0.05. For each contrast extracted with Limma, gene set enrichment of term in the GO Biological Processes 2021 database were detected using web-based EnrichR^52^ by checking the number of significantly differentially expressed genes in each contrast to the known terms and calculating enrichment score and P-value. The volcano plot was generated on the Limma extracted contrasts by plotting the -log_10_(P-value) against the log_2_FC. The R/Bioconductor package heatmap3^80^ was used to display heatmaps for the top 20 differentially expressed genes in each contrast with a Benjamini-Hochberg false-discovery rate adjusted p-value less than or equal to 0.05.

For qRT-PCR, cDNA was synthesized using the SuperScript III First-Strand kit (Invitrogen) using oligo(dT) and used for quantitative PCR in a Real-Time PCR System (Bio-Rad CFX96). Target gene cDNAs was detected using the iTaq Universal SYBR Green master mix (Bio-Rad) and the following primer pairs:

*Gapdh* forward CTTTGTCAAGCTCATTTCCTGG and reverse TCTTGCTCAGTGTCCTTGC^81^, *Oas1* forward CGCACTGGTACCAACTGTGT and reverse CTCCCATACTCCCAGGCATA^54^, *Oas2* forward TTGAAGAGGAATACATGCGGAAG and reverse GGGTCTGCATTACTGGCACTT^54^, *Mx2* forward GAGGCTCTTCAGAATGAGCAAA and reverse CTCTGCGGTCAGTCTCTCT^54^, *Mlkl* forward GGATTGCCCTGAGTTGTTGC and reverse AACCGCAGACAGTCTCTCCA^82^, *Ifnb1* forward CGAGCAGAGATCTTCAGGAAC and reverse TCACTACCAGTCCCAGAGTC^81^, and *Isg15*forward GCAGACTCCTTAATTCCAGGG and reverse TTCAGTTCTGACACCGTCATG^83^. The 2^-ddCt^ method was used to analyze the gene expression fold change after normalization with *Gapdh*.

### Statistical analysis

All data are from at least two independent infection experiments with multiple mice in each group. No blinding was performed during animal experiments. Statistical differences were calculated using Prism 9 (GraphPad) using Mantel-Cox tests and Gehan-Breslow-Wilcoxon test (survival), unpaired two-tailed Mann-Whitney tests (to compare two groups with nonparametric data distribution), and one-way ANOVA with Tukey’s multiple comparisons tests (to compare more than two groups with parametric data distribution). Differences with a p-value of <0.05 were defined as statistically significant. Statistical outcomes for each comparison for all graphs are available in **Table S4**.

## Supporting information

Table S1: List of genes significantly downregulated at 21dpi in the lungs of Irgm1-/-mice compared to WT mice.

Table S2: List of genes significantly upregulated at 21dpi in the lungs of Irgm1-/- mice compared to WT mice.

List of genes significantly upregulated in the lungs of uninfected Irgm1-/- mice compared to WT mice.

Statistical analysis of samples in the graphs from all figures.

## Data availability

RNA-seq data have been deposited in the NCBI Gene Expression Omnibus (GEO) database and are accessible through the GEO SuperSeries accession number GSExxx. All other relevant data are available from the corresponding author upon reasonable request.

## Acknowledgments

This work was supported by NIH grants R01 AI132697 and U19 AI142784, a Burroughs Wellcome Fund Investigators in the Pathogenesis of Infectious Disease Award, and the Philip and Sima Needleman Center for Autophagy Therapeutics and Research to C.L.S., a Stephen I. Morse Fellowship to S.K.N., NIH grant T32 AI007172 to M.E.M., a Potts Memorial Foundation postdoctoral fellowship to R.L.K., and a Alexander & Gertrude Berg Fellowship to N.D. We also acknowledge the Genome Technology Access Center at the McDonnell Genome Institute at Washington University School of Medicine for RNA sequencing and analysis. The Center is partially supported by NCI Cancer Center Support Grant #P30 CA91842 to the Siteman Cancer Center and by ICTS/CTSA Grant# UL1TR002345 from the National Center for Research Resources (NCRR), a component of the National Institutes of Health (NIH), and NIH Roadmap for Medical Research. Schematic diagrams were made with BioRender.com. This publication is solely the responsibility of the authors and does not necessarily represent the official view of NCRR or NIH. We would also like to thank all Stallings lab members for their fruitful suggestions and discussion.

## Author contributions

S.K.N. and C.L.S. conceived the study and designed the experiments. S.K.N., M.E.M., R.L.K., A.S., C.S.C., S.M., N.D. and R.W. performed the experiments. Y.M. analyzed the RNAseq data. D.K. took care of the mouse breeding and mouse-house facility. S.K.N and C.L.S. wrote the manuscript. M.E.M., Y.M., R.L.K., A.S., C.S.C., S.M., N.D. and R.W. edited the manuscript. C.L.S. directed the study, provided funding, and finalized the manuscript.

## SUPPLEMENTAL MATERIAL

**Fig. S1:**
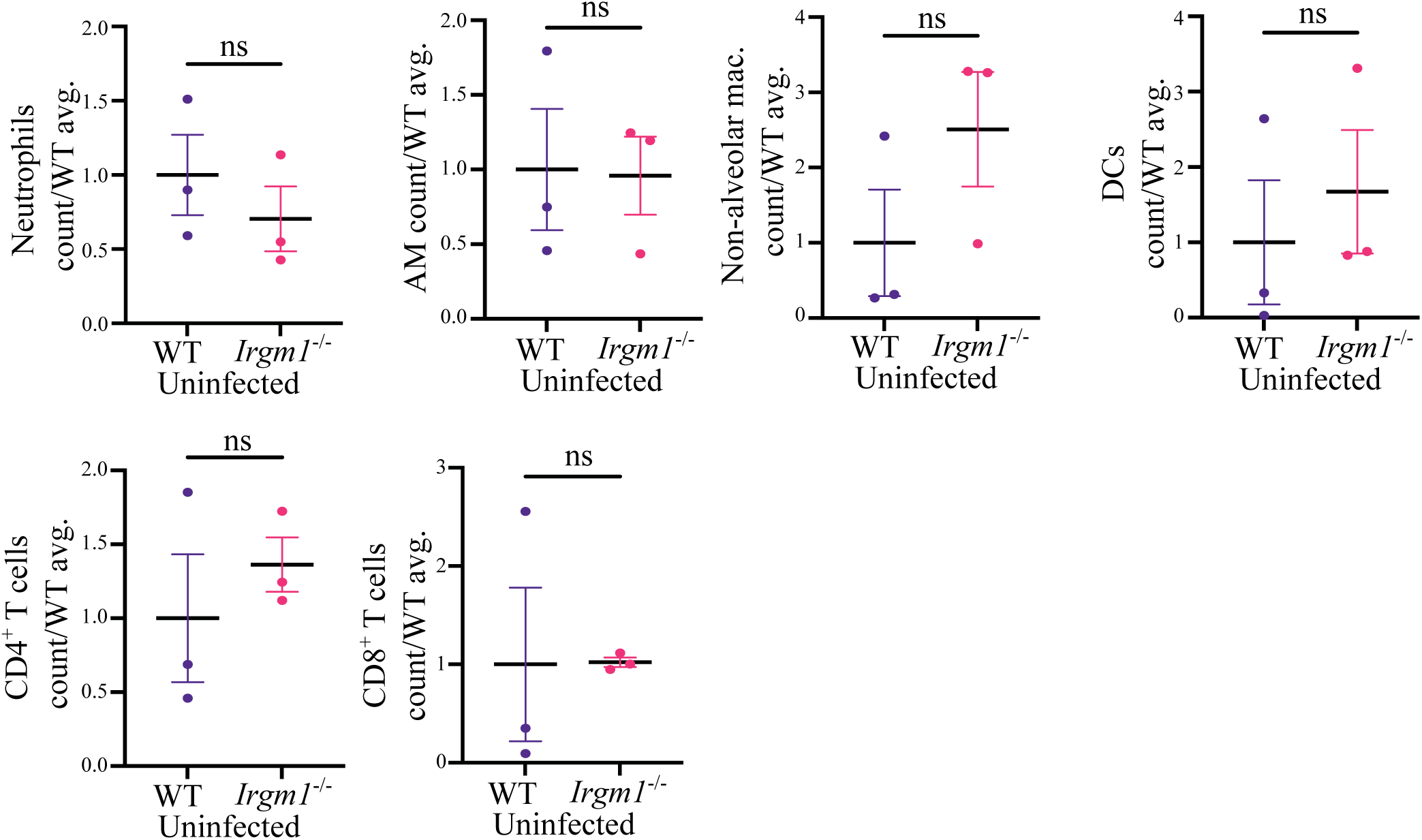
Immune cell populations in the lungs of WT and *Irgm1*^-/-^ mice. Number of neutrophils, AMs, non-alveolar macrophages, DCs, CD4^+^ T cells, and CD8^+^ T cells in the lungs of uninfected WT and *Irgm1*^-/-^ mice. Each data point corresponds to one mouse and graphs show the mean ± SEM, and the data are expressed as the ratio to the average for WT mice. Statistical significance of differences was determined by two-tailed unpaired t-test. ns=not significant.

**Fig. S2:**
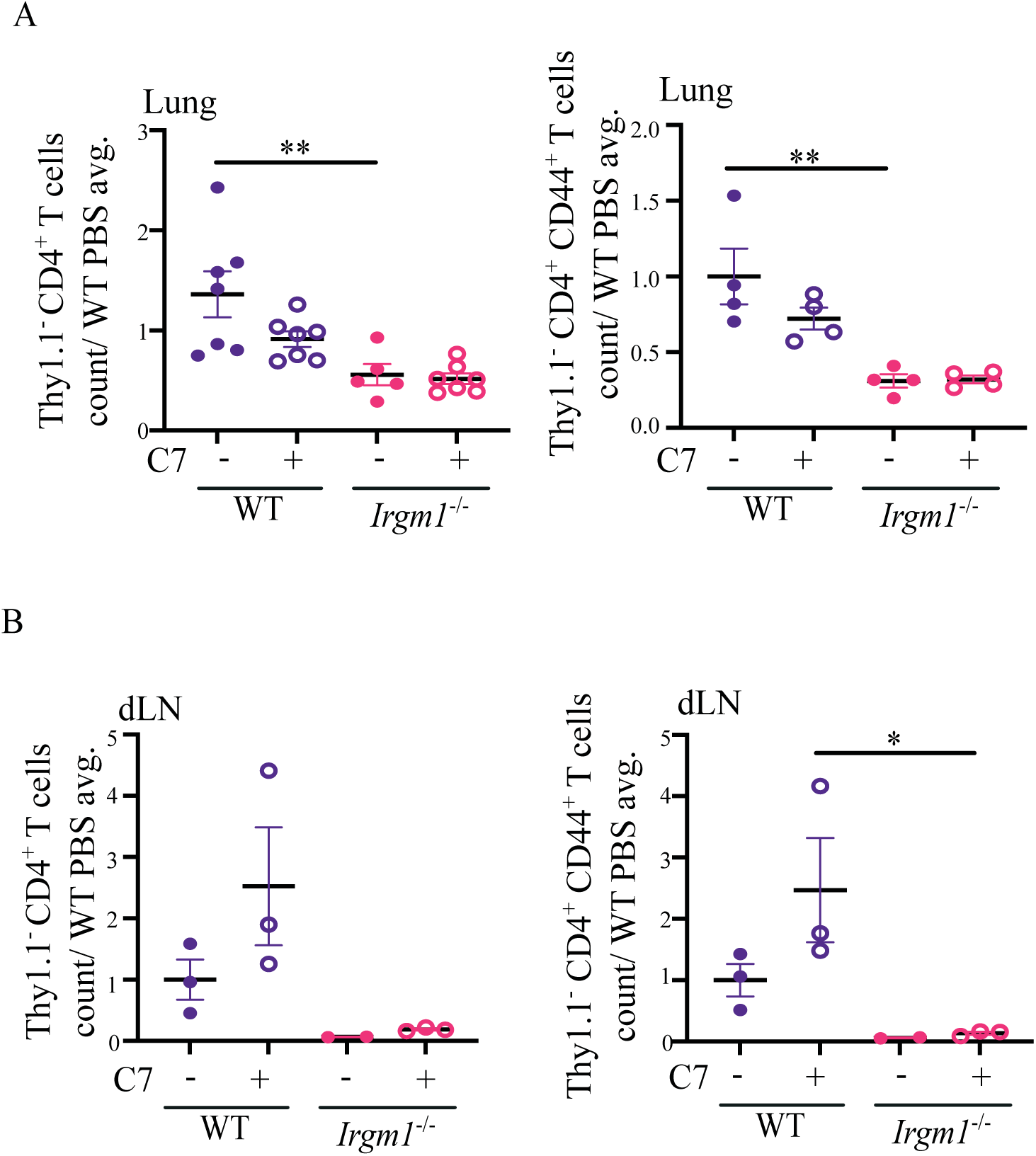
Recipient T cells in adoptive transfer experiments. **(A-B)** Number of the host-derived (Thy1.1^-^) CD4^+^ T cells (left) and activated CD4^+^ T cells (CD44^+^) (right) in the **(A)** lungs and **(B)** dLN at 21 dpi in mice that either received activated C7 CD4^+^ T cells (+) or PBS (-). Each data point corresponds to one mouse and graphs show the mean ± SEM, and the data are expressed as the ratio to the average for WT mice. Statistical significance of differences within a single mouse strain or treatment condition were determined by one-way ANOVA’s Tukey’s multiple comparison test and only significant differences are shown. ** P < 0.01; * P < 0.05.

**Fig. S3:**
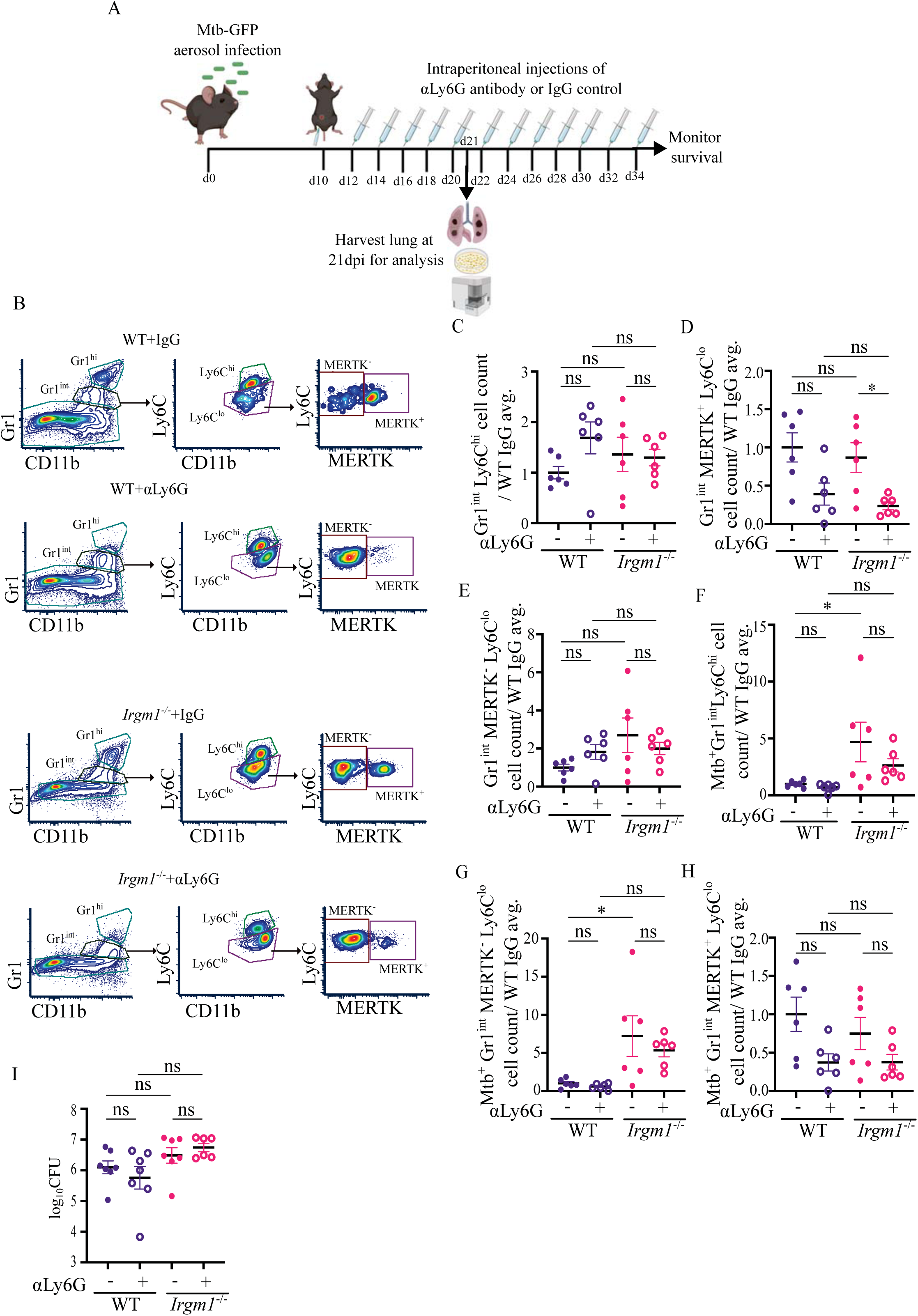
Analysis during neutrophil depletion. **(A)** Schematic of neutrophil depletion in WT and *Irgm1*^-/-^ mice. **(B)** Representative flow-plot showing Gr1^int^ cells with distinct Ly6C^hi^, MERTK^+^Ly6C^lo^, and MERTK^-^Ly6C^lo^ cell populations in the lung at 21dpi. **(C-E)** Number of **(C)** Gr1^int^Ly6C^hi^ **(D)** Gr1^int^MERTK^+^Ly6C^lo^ **(E)** Gr1^int^MERTK^-^Ly6C^lo^ cells in the lungs at 21dpi following administration of αLy6G antibody (+) or an IgG control antibody (-). **(F-H)** Number of Mtb infected **(F)** Gr1^int^Ly6C^hi^ **(G)** Gr1^int^MERTK^-^ Ly6C^lo^ **(H)** Gr1^int^MERTK^+^Ly6C^lo^ cells in the lungs at 21 dpi following administration of αLy6G antibody (+) or an IgG control antibody (-). **(I)** Related to Fig. 3M: Graph showing total increased CFU count in the lung of *Irgm1*^-/-^ mice; the values were statistically not significant because of the variation between experiments. Each data point corresponds to one mouse and the mean ± SEM is shown, where the data is expressed as the ratio to the average for WT mice administered IgG control antibody. Statistical significance of differences within a single mouse strain or treatment condition were determined by one-way ANOVA’s Tukey’s multiple comparison test. ns=not significant.

**Fig. S4:**
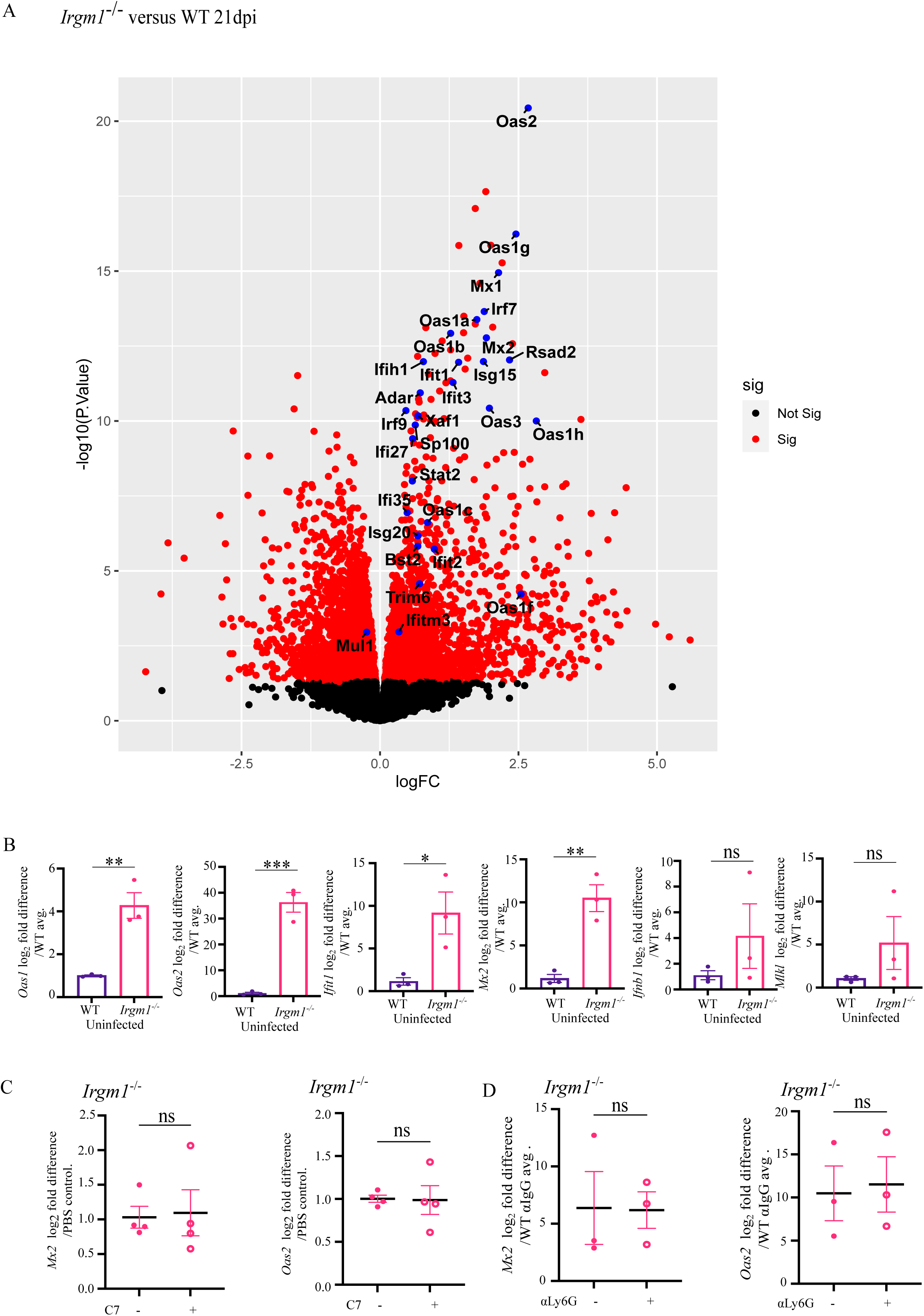
ISG expression in WT and *Irgm1^-/-^* mice. **(A)** Volcano-plot of gene expression in the lungs of *Irgm1^-/-^* versus WT mice at 21 dpi with the significantly differentially expressed genes in red and the genes belonging to the “Cellular Response to Type I IFN pathway” (GO:0071357) in blue. **(B-D**) qRT-PCR analysis of ISG expression in lungs of **(B)** naïve *Irgm1^-/-^* mice compared to WT controls, **(C)** *Irgm1^-/-^* mice at 21 dpi in mice that either received activated C7 CD4^+^ T cells (+) or PBS (-), and **(D)** *Irgm1^-/-^* mice at 21 dpi following administration of αLy6G antibody (+) or an IgG control antibody (-). Each data point corresponds to one mouse and the mean ± SEM is shown, where the data is expressed as the ratio to the average for **(B)** WT mice, **(C)** *Irgm1^-/-^* mice administered PBS, and **(D)** WT mice administered IgG control antibody. Statistical significance of differences within a single mouse strain or treatment condition were determined by Significance of differences were determined by **(B-D)** two-tailed unpaired t-test. *** P < 0.001; ** P < 0.01; * P < 0.05.

**Fig. S5:**
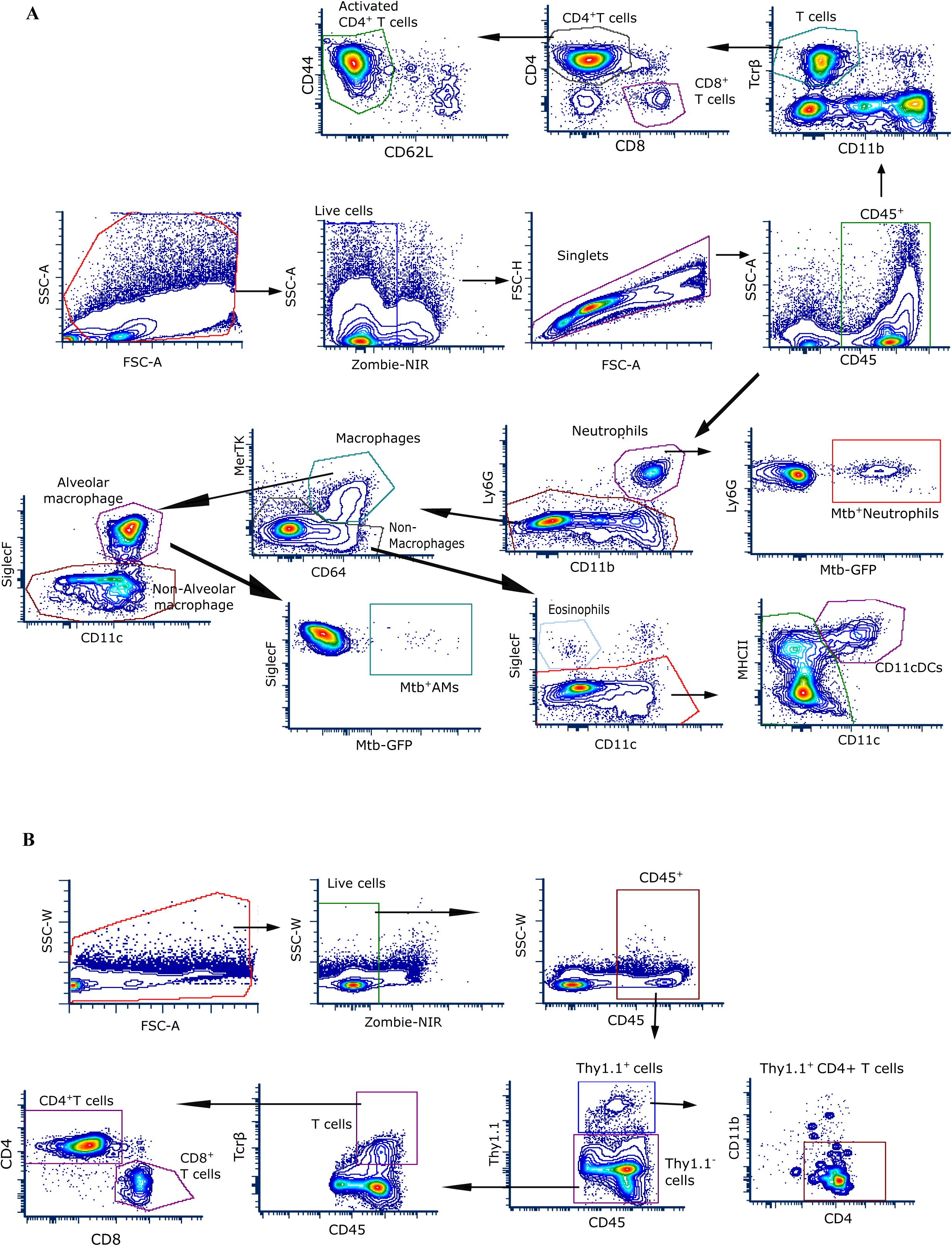
Flow cytometry gating strategy. **(A**) Flow cytometry gating strategy followed for immune cells in murine lungs and dLN. **(B)** Gating strategy for Thy1.1^+^ and Thy1.1^-^ cells for C7 transgenic CD4^+^ T cell adoptive transfer experiment (related to Fig. 2 and Fig. S2). Flow plots were prepared with FCS Express software.

**Table S1.** List of genes significantly (p_adj._ :≤ 0.05) downregulated at 21dpi in the lungs of *Irgm1*^-/-^ mice compared to WT mice.

**Table S2.** List of genes significantly (p_adj._ :≤ 0.05) upregulated at 21dpi in the lungs of *Irgm1*^-/-^ mice compared to WT mice.

**Table S3.** List of genes significantly (p_adj._ :≤ 0.05) upregulated in the lungs of uninfected *Irgm1*^-/-^ mice compared to WT mice.

**Table S4.** Statistical analysis of samples in the graphs from all figures.

